# Accelerating Gene Discovery by Phenotyping Whole-Genome Sequenced Multi-Mutation Strains and Using the Sequence Kernel Association Test (SKAT)

**DOI:** 10.1101/027540

**Authors:** Tiffany A. Timbers, Stephanie J. Garland, Swetha Mohan, Stephane Flibotte, Mark Edgley, Quintin Muncaster, Vinci Au, Erica Li-Leger, Federico I. Rosell, Jerry Cai, Suzanne Rademakers, Gert Jansen, Donald G. Moerman, Michel R. Leroux

## Abstract

Forward genetic screens represent powerful, unbiased approaches to uncover novel components in any biological process. Such screens suffer from a major bottleneck, however, namely the cloning of corresponding genes causing the phenotypic variation. Reverse genetic screens have been employed as a way to circumvent this issue, but can often be limited in scope. Here we demonstrate an innovative approach to gene discovery. Using *C. elegans* as a model system, we used a whole-genome sequenced multi-mutation library, from the Million Mutation Project, together with the Sequence Kernel Association Test (SKAT), to rapidly screen for and identify genes associated with a phenotype of interest, namely defects in dye-filling of ciliated sensory neurons. Such anomalies in dye-filling are often associated with the disruption of cilia, organelles which in humans are implicated in sensory physiology (including vision, smell and hearing), development and disease. Beyond identifying several well characterised dye-filling genes, our approach uncovered three genes not previously linked to ciliated sensory neuron development or function. From these putative novel dye-filling genes, we confirmed the involvement of BGNT–1.1 in ciliated sensory neuron function and morphogenesis. BGNT–1.1 functions at the *trans*-Golgi network of sheath cells (glia) to influence dye-filling and cilium length, in a cell non-autonomous manner. Notably, BGNT–1.1 is the orthologue of human B3GNT1/B4GAT1, a glycosyltransferase associated with Walker-Warburg syndrome (WWS). WWS is a multigenic disorder characterised by muscular dystrophy as well as brain and eye anomalies. Together, our work unveils an effective and innovative approach to gene discovery, and provides the first evidence that B3GNT1-associated Walker-Warburg syndrome may be considered a ciliopathy.

**Author Summary:** Model organisms are useful tools for uncovering new genes involved in a biological process *via* genetic screens. Such an approach is powerful, but suffers from drawbacks that can slow down gene discovery. In forward genetics screens, difficult-to-map phenotypes present daunting challenges, and whole-genome coverage can be equally challenging for reverse genetic screens where typically only a single gene’s function is assayed per strain. Here, we show a different approach which includes positive aspects of forward (high-coverage, randomly-induced mutations) and reverse genetics (prior knowledge of gene disruption) to accelerate gene discovery. We paired a whole-genome sequenced multi-mutation *C. elegans* library with a rare-variant associated test to rapidly identify genes associated with a phenotype of interest: defects in sensory neurons bearing sensory organelles called cilia, *via* a simple dye-filling assay to probe the form and function of these cells. We found two well characterised dye-filling genes and three genes, not previously linked to ciliated sensory neuron development or function, that were associated with dye-filling defects. We reveal that disruption of one of these (BGNT–1.1), whose human orthologue is associated with Walker-Warburg syndrome, results in abrogated uptake of dye and cilia length defects. We believe that our novel approach is useful for any organism with a small genome that can be quickly sequenced and where many mutant strains can be easily isolated and phenotyped, such as *Drosophila* and *Arabidopsis*.

## Introduction

A powerful, tried and true approach to identify which genes function in a particular biological process is to create collections of organisms harbouring multiple mutations *via* random mutagenesis, followed by screening the mutant library for organisms that exhibit the desired altered phenotypes. Although such forward genetics strategies have produced numerous fundamental discoveries, a significant limitation of this approach in metazoans is the prolonged time required to identify the causative mutations. The bottleneck typically arises from the required genetic mapping, complementation tests to exclude known genes, and sequencing of candidates genes.

To circumvent the major disadvantage of forward genetics, reverse genetic approaches have been employed. Various strategies for disrupting a collection of known genes (*e.g.,* RNAi, homologous recombination, transposon mutagenesis, *etc.*) are combined with phenotypic screening to identify candidates. Reverse genetics approaches also have drawbacks, however, including the need to handle and process tens of thousands of strains to assay the entire genome, off-target effects in the case of RNAi, and omission of essential genes.

We hypothesised that we could use whole-genome sequencing in combination with statistical genetics to inaugurate a novel gene discovery approach which retains the advantages of both forward and reverse genetics, and yet minimises their downsides. To do this, we employed the Million Mutation Project (MMP) [1], a collection of 2007 *Caenorhabditis elegans* strains harbouring randomly-induced mutations whose genomes are fully sequenced (data is publicly available: (http://genome.sfu.ca/mmp/about.html). This mutant library represents an unprecedented genetic resource for any multicellular organism, wherein the strains collectively contain one or more potentially disruptive alleles affecting nearly all *C. elegans* coding regions. On average, each strain contains ∼ 400 non-synonymous mutations affecting protein coding sequences.

We postulated that this whole-genome sequence information would allow an “eyes wide open” approach when performing a genetic screen, such that pairing this resource with a high-throughput assay would enable rapid discovery of genes not previously associated with our biological process of interest. Here, we demonstrate that testing for association between variants from the MMP library and phenotype data with the Sequence Kernel Association test (SKAT) [2] allows us to effectively and efficiently predict novel genes important for our chosen biological process: the development and function of the amphid and phasmid sensillum, which includes both ciliated sensory neurons as well as glial-like neuronal support cells.

Primary (non-motile) cilia arise from a modified centriole (basal body) and act as ‘cellular antennae’ that transduce environmental cues to the cell [3]. They enable sensory physiology (such as olfaction/chemosensation, mechanosensation, vision) and are central to signalling pathways essential for metazoan development [4]. Dysfunction of cilia is implicated in a number of human diseases, including polycystic kidney disease, congenital heart disease, and an emerging group of genetic disorders termed ciliopathies (*e.g.*, Bardet-Biedl, Meckel-Gruber and Joubert Syndromes). In these ciliopathies, disruption of many, if not all, cilia in the human body results in a plethora of defects, including retinal degeneration, organ cyst formation, obesity, brain malformations, and various other ailments [5][6].

In *C. elegans,* the uptake of a fluorescent lipophilic dye, Dil, from the environment is used to probe the integrity of the amphid and phasmid sensillum, which includes cilia and ciliated sensory neurons, as well as glial-like sheath cells. Dil is selectively incorporated into six pairs of ciliated amphid channel sensory neurons in the head (ADF, ADL, ASH, ASI, ASJ, and ASK), and two pairs of ciliated phasmid channel sensory neurons in the tail (PHA and PHB), *via* environmentally-exposed cilia present at the tips of dendrites (S1 Fig) [7,8]. Many dye-filling *(dyf)* mutants known from genetic screens [8,9] harbour mutations in genes influencing ciliated sensory neuron development and function, including ciliogenesis [10], cilia maintenance [11], axon guidance [9], dendrite anchoring/formation [10], as well as cell fate [12]. Importantly, noncell autonomous effects from disruption of neural support (glial) cells can also result in dye-filling defects [10,13].

When we applied SKAT to the phenotype data we collected from screening the MMP strains for dye-filling, we found that a previously uncharacterized *C. elegans* gene, *bgnt–1.1/F01D4.9*, plays an essential role in this process. We found that the ciliated sensory neurons of *bgnt–1.1* mutants fail to fill with a lipophilic dye, a phenotype indicative their dysfunction, and that BGNT–1.1 localises specifically to the *trans*-Golgi network of the amphid and phasmid sheath cells. These are glial-like neuronal support cells, which are critical for the development and function of ciliated sensory neurons. Interestingly, *bgnt–1.1* is the orthologue of human B3GNT1/B4GAT1, a gene implicated in Walker-Warburg syndrome [14,15], a disorder with clinical ailments resembling a ciliary disease (ciliopathy).

## Results

We screened 480 randomly-chosen whole-genome sequenced multi-mutation MMP strains, ∼25% of the library, for defects in DiI uptake in amphid and phasmid ciliated sensory neurons (Fig 1). We found 40 MMP strains which exhibit significant amphid dye-filling defects and 40 MMP strains which exhibit significant phasmid dye-filling defects; the strains with amphid and phasmid dye-filling defects are not necessarily identical (Fig 1C, Table 1, **S1 Table**).

**Table 1.** Summary of dye-fill phenotype classes observed

| Phenotype summary | Number of strains |
| --- | --- |
| Amphid and phasmid dye-fill defect | 11 |
| Amphid and phasmid partial dye-fill defect | 12 |
| Phasmid only partial dye-fill defect | 17 |
| Amphid only partial dye-fill defect | 17 |
| Wild-type dye-filling | 423 |

**Fig 1.**
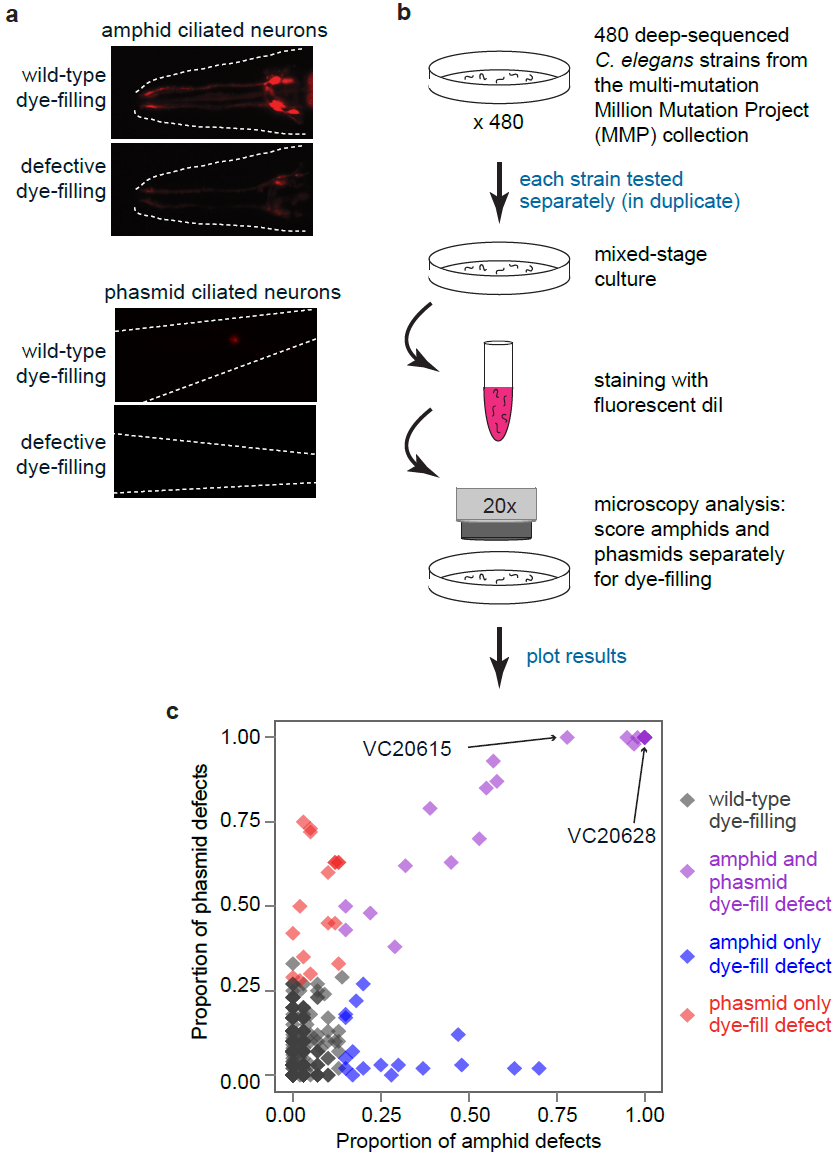
**Dye-filling (ciliated sensory neuron development/function) screening methodology and results.** (A, B) Input to the screen was 480 whole genome-sequenced multi-mutant strains from the Million Mutation Project [10]. Mixed-stage *C. elegans* cultures were incubated with DiI for 30 minutes, washed in buffer and then examined by fluorescence microscopy for their ability to uptake the dye into head (amphid) and tail (phasmid) sensory neurons. (C) Dye-filling phenotypes of each of the 480 MMP strains which were assayed. The proportion of worms exhibiting dye-filling in strains represented by dark grey diamonds were not statistically separable from the proportion of wild-type worms exhibiting dye-filling as assessed by a Fisher’s exact test with p-values adjusted for a 5% false discovery rate (Benjamini-Hochberg procedure) to control for multiple testing. Blue and red diamonds represent strains with mainly or exclusively amphid or phasmid dye-filling defects, while purple diamonds show strains with defects in both amphid and phasmid sensory neurons. Two highlighted strains, VC20615 and VC20628, contain mutations which alter conserved amino acid residues in the protein encoded by *C. elegans bgnt–1.1,* a gene identified by SKAT to be associated with both amphid and phasmid dye-filling defects.

We identified 11 completely dye-fill defective strains, where all worms sampled failed to take up dye. A preliminary look at the data indicates that of these, 10 contained deleterious (“knockout”) mutations in previously identified dye-filling genes (*e.g.*, nonsense and frameshift-inducing deletions; **S2 Table**). Additionally, we uncovered 47 partially dye-fill defective strains, where a proportion of the population failed to fill with dye significantly more often than wild-type worms. Of these partially dye-fill defective strains, 1 harbours a nonsense mutation and 10 display missense mutations in known dye-filling genes (**S2 Table**).

Despite the fact that we can identify some strains with mutations in genes previously shown to cause dye-filling defects, it is not clear that it is the mutations in these genes which are necessarily the cause of the dye-filling defects in these strains. Furthermore, there are 38 strains where we cannot even generate a hypothesis as to what genetic variation is responsible for the dye-filling defect. To facilitate identification of genes responsible for the observed dye-fill defects, we hypothesised that a recently developed statistical genetics approach commonly used in human genetics, but under-utilised in model organisms, would allow for the rapid prioritisation of candidate genes. Specifically we chose to employ the sequence kernel association test (SKAT) to uncover genes associated with the dye-filling phenotype. SKAT is a regression method to test for association between rare and/or common genetic variants in a region and a continuous or dichotomous trait [2].

We chose SKAT over other statistical analyses for several reasons. For our dataset, it was imperative that we chose an association test that dealt effectively with rare variants, as 800,000/850,000 of the non-synonymous variants in the MMP library are unique; meaning that they are present in only a single isogenic strain in the library. Hence, genome-wide association study (GWAS) approaches, which typically test for an association between common variants (generally defined as a minor allele frequency > 5%) and a trait of interest, would be unsuitable for analysis of phenotype datasets derived from the MMP library. We also viewed SKAT as an optimal method to use for our dataset because it permits the use of prior information to assign weights to genetic variants. For example, nonsense mutations might be expected to be more deleterious than other variants which may cause more modest changes to the protein, such as missense mutations and in-frame deletions. The C-alpha test [16], which is quite similar to SKAT in the absence of covariants (*e.g.,* age, sex, *etc.*), could have also been used for our dataset, but we chose to employ SKAT because it facilitates implementing and assigning biologically relevant weights to variants. Finally, SKAT was chosen over other related burden tests, such as the cohort allelic sums test (CAST) [17] and the combined multivariate and collapsing (CMC) method [18], because unlike these tests, SKAT does not assume that all (common) variants will affect the trait in the same direction.

Given that the groups of worms which have amphid dye-filling and phasmid dye-filling defects do not necessarily overlap, we performed SKAT separately for each dataset. We chose to perform the linear regression version of SKAT in combination with log transformation of the response (phenotype) variable, as opposed to a logistic regression version of SKAT because in its current implementation, the logistic regression version of SKAT does not work with proportion data, and takes only dichotomous traits coded as 0 or 1. Quantile-quantile (QQ)-plots were used to choose the appropriate constant to add to the response (phenotype) variable before log transformation (S2 & S3 Fig). Finally, we performed SKAT with biologically relevant weights assigned to the variants. We assigned mutations which would likely result in the creation of a null mutation (nonsense and splicing mutations, as well as frameshift causing deletions) a weight of 1, mutations which would result in truncation of the protein (in-frame deletions) a weight of 0.75, and mutations which would result in a change in amino acid sequence (missense mutation) a weight of 0.25. We hypothesised these were reasonable weights to assign to each class of mutation based on the current knowledge in the field of genetics.

Genome-wide SKAT analyses using biologically relevant weights on the amphid dye-filling dataset reveal 5 genes that reach significance when we adjust for multiple testing using a false discovery rate (FDR; Benjamini-Hochberg procedure) of 5% (FDR adjusted p-value was < 0.05, Table 2, **S3 Table**). SKAT analyses using biologically relevant weights on the phasmid dye-filling dataset uncovered 3 genes which reached significance, again using a FDR of 5% (FDR adjusted p-value was < 0.05, Table 3, **S4 Table**). Dye-filling defects of both amphid and phasmid ciliated neurons is significantly associated with genes encoding intraflagellar transport proteins (OSM–1 and CHE–3), and a glycosyltransferase (BGNT–1.1; Table 2 and 3). Amphid-specific dye-filling defects are found to be associated with genes encoding an Arf-GAP related protein, CNT–1, as well as a mitotic spindle assembly checkpoint protein, MDF–1 (Table 2). No gene was found to be significantly associated with only phasmid dye-filling defects (Table 2 and Table 3).

**Table 2.**
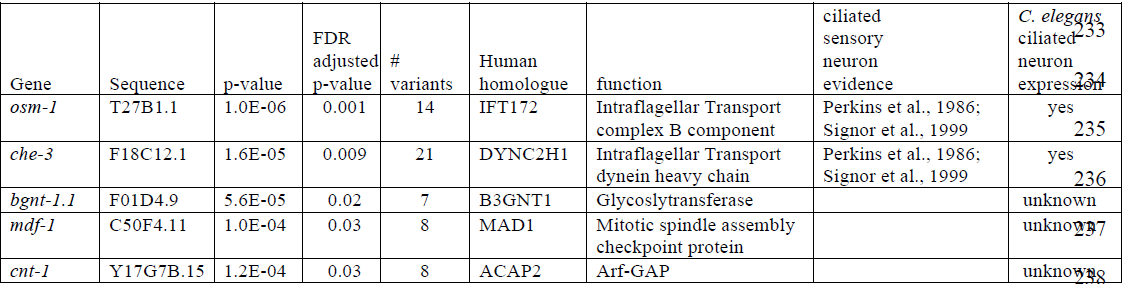
Genes with genome-wide significance for amphid ciliated neuron dye-filling phenotypes, ordered by FDR adjusted p-value.

**Table 3.**
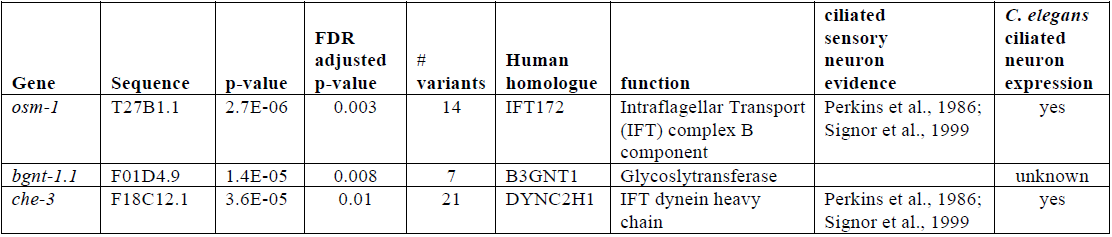
Genes with genome-wide significance for phasmid ciliated neuron dye-filling phenotypes, ordered by FDR adjusted p-value.

Of the three genes associated with both amphid and phasmid dye-filling defects, namely *osm*–1, *che*–3, and *bgnt*–1.1, the first two are well characterised genes whose dye-filling defective phenotypes are ascribed to their key roles in intraflagellar transport (IFT). OSM–1 is the orthologue of mammalian IFT172, an IFT-B subcomplex component which functions as an adaptor to link ciliary cargo (*e.g.,* tubulin, receptors and signaling molecules) to the anterograde IFT kinesin motors, and is necessary for ciliogenesis [10]. CHE–3, the orthologue of mammalian DYNC2H1, is a cytoplasmic dynein heavy chain which powers the retrograde IFT-dynein motor. This molecular motor recycles IFT machinery from the growing ciliary tip back to the ciliary base and is also necessary for proper cilium formation/maintenance [19,20]. These two known dye-fill/cilia genes represent excellent positive controls for our screen, and indicate that other genes found to be significantly associated with these phenotypes may be novel dye-fill genes that influence cilia function.

Interestingly, one of the other amphid dye-filling gene hits, *cnt*–1, encodes a protein that play roles in membrane trafficking/dynamics by influencing small GTPase function, *via* GTPase-activating protein or GAP activity. The general involvement of small GTPases of the Arf, Arf-like (Arl) and Rab families in cilium formation/development is well established [3]. *cnt–1* encodes the orthologue of human ACAP2, which interacts with both Rab35 [21] and Arf6 [22] to mediate crosstalk between these two proteins, at least in the context of PC12 cell neurite outgrowth, and potentially through endocytic recycling [23]. Another amphid dye-filling gene hit, *mdf-1,* is homologous to *Mad1*, and encodes a Mitotic spindle assembly checkpoint protein [24]. To the best of our knowledge, our findings are the first to directly implicate *mdf-1/Mad1* as being important for cilia development and/or function but other Mitotic spindle assembly checkpoint proteins have previously been linked to cilia, including BUBR1 [25] and APC-Cdc20 [26].

The third, putative novel dye-filling gene significantly associated with both amphid and phasmid dye-fill phenotypes is *bgnt–1.1* (Table 2 & 3, **S3** and **S4 Table**). *bgnt–1.1* encodes an unstudied *C. elegans* glycosyltransferase 49 family member homologous to human B3GNT1/B4GAT1 (S4 Fig). B3GNT1 catalyses the addition of P1–3 linked N-acetylglucosamine to galactose [27]. In HeLa cells, its subcellular localisation is concentrated at the *trans*-Golgi [28]. B3gnt1 knockout mice exhibit axon guidance phenotypes [29,30] and deficient behavioural responses to estrous females [31]. In humans, mutations in B3GNT1 are associated with a congenital muscular dystrophy with brain and eye anomalies, Walker-Warburg syndrome (WWS) [14,15]. WWS is a suspected, but unconfirmed ciliopathy; it exhibits 6 core features common to ciliopathies, including Dandy-Walker malformation, hypoplasia of the corpus callosum, mental retardation, posterior encephalocele, retinitis pigmentosa and *situs inversus* [5]. Additionally, one patient is reported to exhibit dysplastic kidneys [14], a developmental disruption which leads to cyst formation, illuminating a potential 7th core ciliopathy feature to this disorder, renal cystic disease. To divulge a potential connection between B3GNT1 and cilia and/or ciliated sensory neuron function, we sought to confirm the role of *C. elegans* BGNT–1.1 in dye-filling, and analyse its involvement role in ciliated sensory neuron development.

Of the eight MMP strains harbouring mutations in *bgnt–1.1*, three (VC20615, VC20628 and VC20326) exhibited severe dye-fill phenotypes (**S1 Table**). The C -> T missense mutation in *bgnt–1.1* in VC20615 corresponds to P194S alteration in the protein sequence, while VC20628 and VC20326 each harbour an identical G -> A missense mutation in *bgnt–1.1*, which leads to a G205E amino acid change in the protein sequence. Both of these mutations alter conserved amino acid residues (S5 Fig). In the screen we encountered 5 additional strains that harboured missense mutations in *bgnt–1.1* but did not exhibit dye-filling defects. Close examination of the predicted effects of these missense mutations on the amino acid sequence of the protein indicates that these alleles do not lead to amino acid changes in conserved residues (S5 Fig), and thus it is not surprising that these strains do not exhibit dye-fill defects.

To confirm that the mutations in *bgnt–1.1* was responsible for the dye-filling phenotypes in *bgnt–1.1* mutants, we rescued the dye-fill defects by expressing a fosmid containing a wild-type copy of *bgnt–1.1* in an extrachromosomal array (Fig 2). Another way to confirm that disruption of *bgnt–1.1* causes dye-fill defects would be to observe this phenotype in a strain harbouring a knock-out mutation in *bgnt–1.1.* Although there are 49 *bgnt–1.1* alleles available, a knock-out allele of *bgnt–1.1* did not yet exist. There are two insertion/deletion alleles available, *gk1221* and *tm4314*, but both fall within introns and thus unlikely to affect protein function. Thus, we also tested the causality of *bgnt–1.1 via* a relatively efficient SNP mapping approach. We established that the dye-fill phenotypes from VC20615 and VC20628 strains map to the *bgnt–1.1* locus, on chromosome IV between −5 cM and 8 cM (S6 Fig). Notably, in both VC20615 and VC20628 strains, *bgnt–1.1* was the only gene in this region harbouring a mutation which was common to both of these strains.

**Fig 2.**
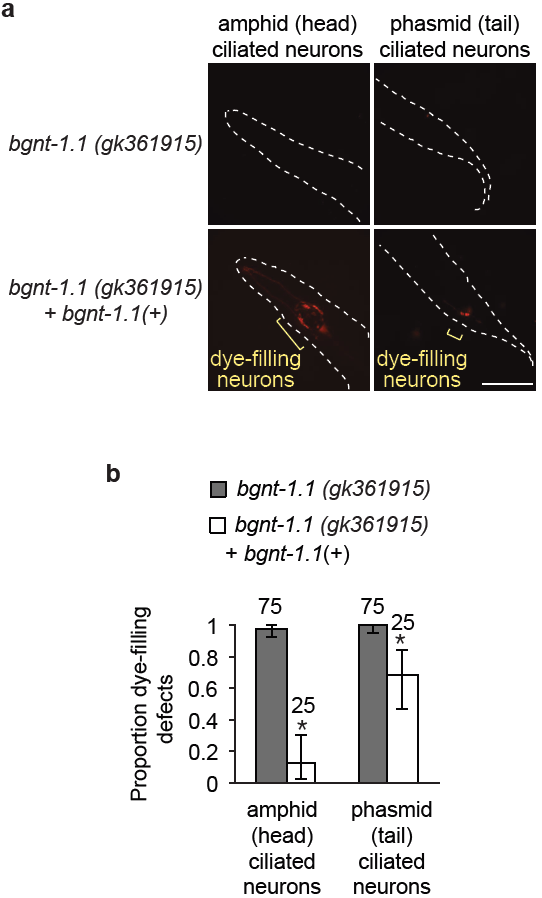
**Confirmation of *bgnt–1.1* as a novel dye-filling gene.** (A) Wild-type *bgnt–1.1* rescues dye-filling defects in *bgnt–1.1 (gk361915)* mutants. Transformation of VC20628 *bgnt–1.1 (gk361915)* with a fosmid containing wild-type *bgnt–1.1* (WRM065bB05) completely rescues the amphid, and partially rescues the phasmid dye-filling defects of VC20628 *bgnt–1.1 (gk361915)* mutants, as assessed by fluorescence microscopy. (B) Quantitation of amphid and phasmid dye-filling in the mutant strain VC20628 in the presence or absence of a fosmid rescue contruct (p < 0.05, Fisher's exact test). Error bars represent 95% confidence intervals (Pearson Clopper method).

Finally, we used CRISPR-Cas9 genome engineering [32] to independently generate three *bgnt–1.1* knockout alleles. All three alleles delete the first and second exon of *bgnt–1.1* and insert a selectable marker, *Pmyo–2::GFP*, in their place. When tested for dye-filling defects, we observe an identical dye-filling phenotype as observed in the 6X outcrossed *bgnt–1.1* (*gk210889*) G205E allele from the Million Mutation Project (Fig 3). In all of these mutants, we observe a great decrease in the amount of dye that enters the amphid ciliated sensory neurons, which is often undetectable, as well as a complete absence of dye-filling of the phasmid ciliated sensory neurons. Together, these findings indicate that loss of *bgnt–1.1* function results in dye-filling defects.

**Fig 3.**
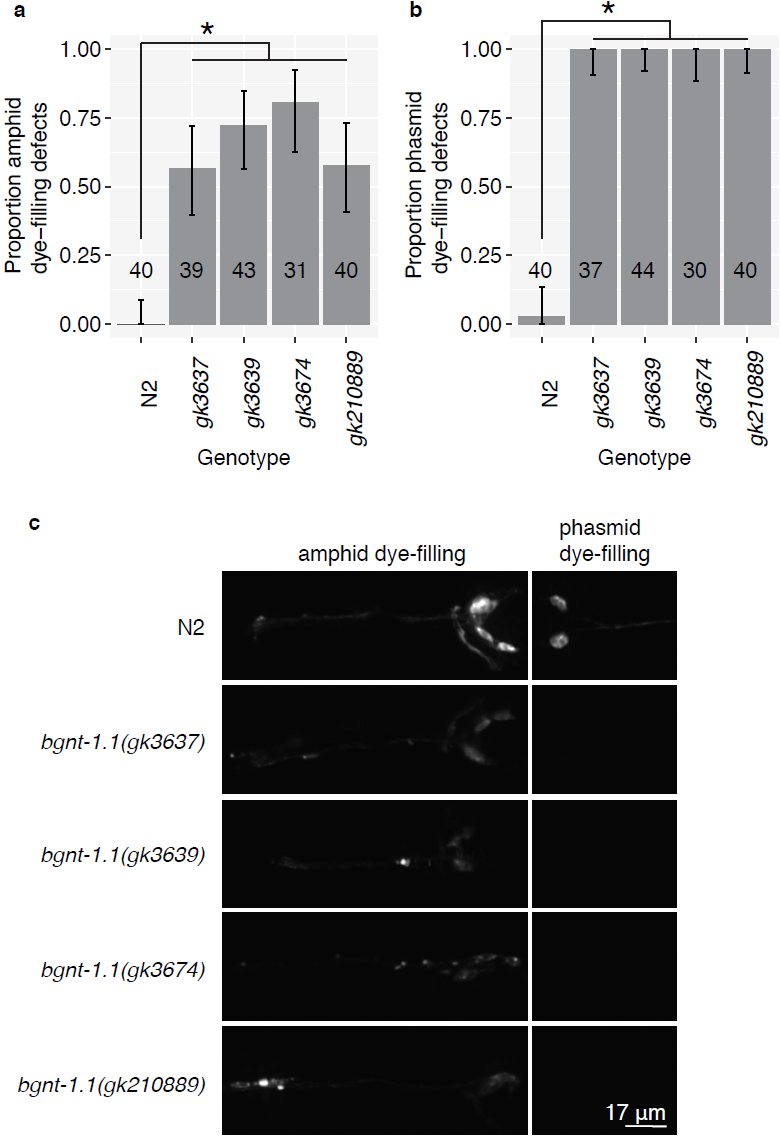
**CRISPR-Cas9 knockout of *bgnt–1.1* causes dye-filling defects.** (A) three *bgnt–1.1* knockout alleles, independently generated via CRISPR-Cas9, exhibit amphid and phasmid (B) dye-fill defects compared to wild-type worms (p < 0.0001, Fisher’s exact test, p-values adjusted for a 5% false discovery rate using the Benjamini-Hochberg procedure). The magnitude of the dye-filling defects in the CRISPR-Cas9 knockout alleles is similar to that observed in the MMP *bgnt–1.1 gk210889* missense mutation, which results in G205E amino acid change. Error bars represent 95% confidence intervals (Pearson Clopper method). (C) Representative images of dye-filling in wild-type (N2), three CRISPR-Cas9-generated *bgnt–1.1* knockout alleles and the MMP *bgnt–1.1 gk210889* missense mutation allele.

To shed light on how *bgnt–1.1* affects dye-filling, we studied expression of GFP-tagged BGNT–1.1 constructed *via* fosmid recombineering (https://transgeneome.mpi-cbg.de/transgeneomics/index.html), and thus containing all of this gene's endogenous regulatory elements. We observed that in *C. elegans*, the protein localises to discrete structures in the cell body of the amphid and phasmid glial-like sheath cells (AMsh and PHsh, respectively) in the head and tail of the animal (Fig 4a). These cells are intimately associated with the ciliated sensory neurons in the pore region where cilia are exposed to the external environment (Fig S1).

**Fig 4.**
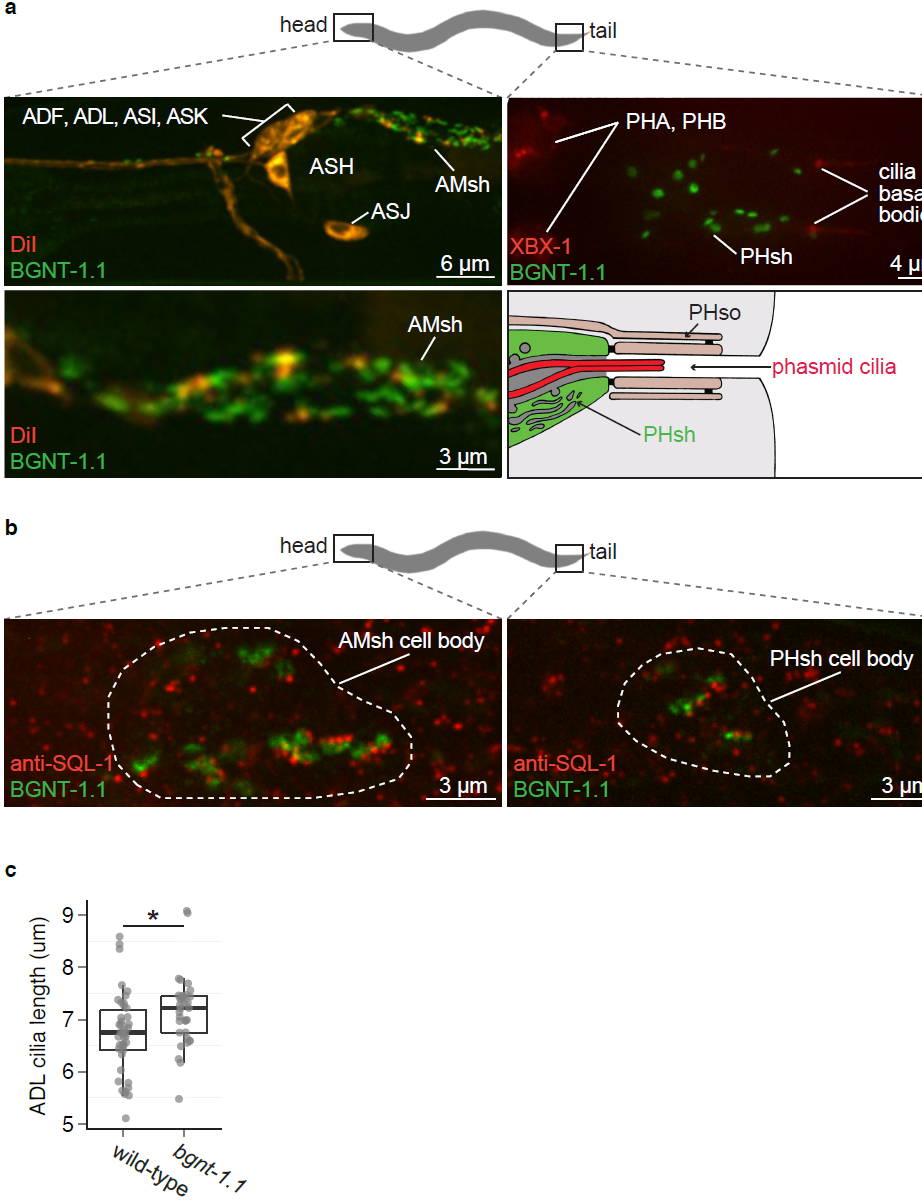
**BGNT-1.1 localises to the *trans*-golgi in the amphid and phasmid sheath cells. (A)** BGNT–1.1::GFP generated from a recombineered fosmid localises to the amphid and phasmid sheath cells. Amphid ciliated neurons (ADF, ADL, ASH, ASI, ASJ, and ASK) were visualized with DiI, while phasmid ciliated neurons (PHA and PHB) were visualized with Posm–5::XBX–1::tdTomato. Amphid sheath cells are abbreviated to AMsh, and phasmid shearh cells are abbreviated to PHsh. (B) BGNT–1.1::GFP localises proximal to the discrete anti-SQL–1 puncta. SQL–1 is an established cis-Golgi marker, indicating that BGNT–1.1 is concentrated at the *trans*-Golgi. (C) ADL cilia are significantly longer in *bgnt–1.1* mutants compared to wild-type (p < 0. 01, Kruskal-Wallis test). ADL cilia are labelled with Psrh–220::IFT–20::tdTomato. IFT–20 (IFT20) localises to cilia basal bodies (bb) and axonemes. The *srh–220* promoter drives expression primarily in ADL neurons.

Mammalian B3GNT1 is found at the *trans*-Golgi network in HeLa cells [28]. To assess whether this is also where *C. elegans* BGNT–1.1 localises, we performed antibody staining for SQL–1, an established cis-Golgi marker [33], in the strain expressing BGNT–1.1::GFP. We observe that in both the head and tail, the localisation of BGNT–1.1::GFP is always proximal to the discrete SQL–1 puncta, indicating that *C. elegans* BGNT–1.1 is concentrated at the *trans*-Golgi, as expected (Fig 4b).

Next, we queried whether loss of *bgnt–1.1* function in the amphid and phasmid sheath cells leads to any gross ciliary morphology defects by expressing a ciliary marker in *bgnt–1.1* mutants, namely the GFP-tagged IFT-B subcomplex protein, CHE–2 (IFT80). This experiment indicates that although the cilia of *bgnt–1.1* mutants fail to fill with dye, their ciliary structures appear superficially wild-type (S7A Fig). Since modest cilia structure defects may be more difficult to observe using pan-cilia markers, due to overlapping ciliary signals, we also characterised the phenotype of cilia and dendrites in *bgnt–1.1* mutants within a single ciliated amphid cell, the ADL neuron. For this purpose, we used the primarily cell-specific ADL promoter, *Psrh–220*, to drive expression of another cilia marker, IFT–20 (IFT20) tagged with tdTomato. In this strain, we also expressed cytoplasmic GFP in the amphid socket cells so that we could evaluate whether or not the ADL cilia were correctly associated with the surrounding glial support cells and the pore where DiI has access to the amphid ciliated sensory neurons from the environment. Similar to the experiment with the CHE–2::GFP pan-cilia marker, the *Psrh*–220::IFT–20::tdTomato marker revealed that the ADL cilia and amphid socket (Amso) cell morphology also appear superficially wild-type in *bgnt–1.1* mutants (S7B Fig). We then sought to assay for potential phenotypes involving ADL cilia length (Fig 4c); length of socket cell penetration by ADL (proxied by the distance from the distal tip of ADL cilia to the distal end of the socket cell tip; S7C Fig); ADL guidance (proportion of double rod cilia/amphid; S7D Fig); and finally, ADL dendrite blebbing (structural alteration where dendrites take bead on a string appearance; S7E Fig). Our analyses reveal that ADL cilia in *bgnt–1.1* mutants are wild-type in most aspects except for a modest cilia length defect. Specifically, *bgnt–1.1* mutants were observed to have significantly longer cilia compared to wild-type worms (Fig 4c; p < 0.01, Kruskal-Wallis test).

BGNT–1.1 therefore influences amphid and phasmid neuron development and function, as well all modestly affects cilium length, without overtly affecting the gross structure of neurons or cilium formation. The localisation of BGNT–1.1 at the *trans*-Golgi network of sheath cells signifies that its effect on ciliated sensory neurons is non-cell autonomous. Interestingly, when the *bgnt–1.1* amphid ciliated sensory neurons do fill with dye, we observe bright accumulations of dye along and/or beside the dendrites (Fig 3c). These are often brighter than the staining of the cell bodies. Accumulations of dye have been observed in wild-type worms and have been attributed to the dye-filling of the amphid sheath cells [34], but these are qualitatively much smaller than what we observed in the *bgnt–1.1* mutants. This suggests a potential alteration in the ability of the sheath cell to take up, or intracellularly distribute dye when BGNT–1.1 is disrupted.

In humans, mutations in *B3GNT1* cause Walker-Warburg syndrome. Given that mutations in B3GNT1 lead to WWS and that it is classified as a dystroglycanopathy, a group of muscular disorders whose etiology is hypothesised to be caused by aberrant glycosylation of dystroglycan, we tested whether or not the *C. elegans* dystroglycan homologs, *dgn-1*, *dgn-2* and *dgn-3*, exhibited dye-filling phenotypes. We observed that all *dgn* mutants exhibited dye-filling indistinguishable from wild-type worms (S9 Fig), indicating that BGNT-1.1 function in dye-filling is likely independent of dystroglycan. Interestingly, as highlighted earlier, the WWS congenital muscular dystrophy exhibits 6 features beyond muscle structure/function disruption which are core ciliary disorder (ciliopathies) features [5]. Our findings that *C. elegans bgnt–1.1* is expressed in glial cells directly associated with, and necessary for the function of ciliated sensory neurons, is consistent with its role in cilium-dependent dye-filling.

## Discussion

Here we demonstrate that rare-variant association analysis (*e.g.*, SKAT) is an efficient way to rapidly uncover novel genes for a phenotype of interest (*e.g.*, ciliated sensory neuron function) in whole-genome sequenced strains harbouring multiple mutations induced *via* random mutagenesis. We found that three cilia-related genes, *osm–1*, *che–3* and *bgnt–1.1* were significantly associated with dye-filling defects, suggesting that the remaining unstudied genes, *cnt–1* and *mdf–1*, likely represent genes important for ciliary/sensory neuron development and/or function.

### The Glycosyltransferase BGNT-1.1 is Essential for Ciliated Sensory Neuron Dye-filling and Cilium Length Control

We confirmed that *bgnt–1.1*, a gene identified by SKAT as being associated with the dye-filling phenotypes but not previously implicated in cilia or amphid-sensillum function, is a *bona fide* dye-filling gene. We observed that: *(1)* two missense mutations in *bgnt–1.1* result in severe dye-fill defects in three MMP strains, and three CRISP-Cas9-mediated *bgnt–1.1* gene disruptions also cause dye-fill defects; *(2)* a fosmid containing full-length wild-type *bgnt–1.1* rescues the dye-filling phenotype in *bgnt–1.1* mutants; *(3)* the dye-filling phenotypes in the MMP strains with mutations in *bgnt–1.1* map to the *bgnt–1.1* locus; *(4)* BGTN-1 is expressed in sheath cells, which are directly implicated in dye-filling; and finally; and *(5)* mutations in *bgnt–1.1* result in a small but statistically significant ciliary length defect. Together, these data strongly indicate that BGNT–1.1, which we find localises as expected to the *trans*-Golgi network, functions in sheath (glia-like) cells to influence dye-filling.

### Considerations of our Novel Gene Discovery Approach

The power of genome-wide rare-variant association analysis (*e.g.*, SKAT) augments as the number of strains increases (the probability of additional mutations in specific genes is increased), and thus, screening the entire MMP library would likely uncover many additional genes associated with dye-filling defects. To try to assess the minimal number of strains that should be assayed with this approach we performed a power analysis. Raw amphid dye-filling phenotype and genotype data was randomly sub-sampled (without replacement) and analysis was performed via SKAT with, and without, biologically relevant weights. This was done 100 times for each sample size (50, 100, 200, 300, 400). For each sample size, power was calculated as the proportion of times the analysis found at least one gene to be significantly associated with the phenotype. We observed that there was 40% power to detect a single gene as being associated with the amphid dye-filling phenotype at N = 400 for both SKAT with and without biologically relevant weights (S9 Fig). Thus, we recommend that future studies using this method should use a sample size close to what was used in this present study (∼ 500) to maximize the probability that a gene(s) will be found that is significantly associated with the phenotype of interest.

We performed SKAT analyses *via* two methods, 1) while applying biologically relevant weights to the variants (**S3** & **S4 Tables**), and 2) while weighting all variants equally (**S5** & **S6 Tables**). SKAT analysis of the 480 strains without weights was less powerful, and resulted in identifying only 3 genes as being significantly associated with the amphid dye-filling phenotype and 1 gene as being significantly associated with the phasmid dye-filling phenotype; compared to 5 genes and 3 genes, respectively, when biologically relevant weights were used. Although there appears to be no difference in power when SKAT is performed with or without weights at smaller sample sizes (S9 Fig). Thus, to maximize the ability to detect genes associated with the phenotype of interest, in addition to recommending a minmum sample size of ∼ 500, we also recommend assigning biologically relevant weights when using SKAT with the MMP library. The weight assignment could be simple, as done here, or more complex, calculating, for example, the SIFT [35] or Polyphen–2 [36] scores for assessing the severity of each variant in the MMP library.

The genome-wide statistical genetic approach presented here has several advantages over traditional screening approaches. It generates a prioritised list of candidate genes likely responsible for the phenotype of interest. After this list is generated *via* screening and SKAT analysis, candidates can be tested for their causality of the phenotype through several standard genetic approaches in *C. elegans.* Candidates could be confirmed, for example, by *(i)* testing for the phenotype in knock-out mutants or RNAi; *(ii)* genetic rescue experiments; *(iii)* performing a genetic complementation test between two loss of function alleles; or, *(iv)* mapping the mutation to the gene locus. This strategy may work for phenotypes where the traditional polymorphic SNP-mapping strain, CB4856, diverges from the reference wild-type strain, N2, from which the MMP library was generated [1], as well as partially-penetrant or other difficult-to-score phenotypes. In the case of *bgnt–1.1*, we performed genetic rescue experiments, SNP mapping and created CRISPR-Cas9 knockout strains to support our the SKAT findings, which together confirm that *bgnt–1.1* mutations cause dye-filling defects.

Another potential extension and utility of this approach that could work for some (non-neural) phenotypes would be pairing the screening of the MMP strains with RNAi to look for enhancing, suppressing or synthetic phenotypes, and then using SKAT to prioritise a list of candidate genes. Furthermore, as more data from multiple phenotypes are collected on the MMP strains, these could be combined to perform multi-variate genome-wide statistical analysis on whole-genome sequence data. Such approaches are more powerful than univariate approaches in the case of SNP array data [37–39], and such tests can also indicate which variants are pleiotropic, or specific to a single phenotype. How to perform this multivariate phenotype analysis on whole-genome sequences is currently an active area of research and tools to make this possible are being developed, with [40] looking promising.

There are also challenges and limitations to the statistical genetic approach presented here. First, this approach of performing a “medium”-scale screen of the MMP strains is limited to assays that can be done without genetic manipulation of the strains. For example, introducing a functional ‘reporter’ (transgene) into 480 strains would require a prohibitive amount of work, although this has been done for 90 MMP strains [41]. Second, the statistical analysis presented here is only possible for genes which have > 1 variant in the population of worms screened. In practice, we found it works optimally for strains with at least 7 variants. This is due to the distribution of p-values when attempting to control for multiple testing; in our dataset, fewer than 7 variants led to a skewed p-value distribution and an inflation of False-discovery rate adjusted p-values. This strict rule demanding high-coverage for our SKAT analysis leads to only 1150 genes in the 480 MMP strains being considered here. This is due to the distribution of variant counts per gene in the MMP strains (S9 Fig), which exponentially decreases from 1 to N.

### Mechanism of BGNT-1.1 in Dye-Filling

How disrupting BGNT–1.1 abrogates dye filling remains uncertain. One possibility is that the glycosyltransferase regulates the association of cilia with the sheath and socket glial-like cells which envelop them (S1 Fig). Specifically, we hypothesise that BGNT–1.1 functions in the *trans*-golgi network of the amphid and phasmid sheath to glycosylate key unidentified proteins important for the association of this sensillum organ. This defect will not be visible at the level of light microscopy, and could perhaps result from changes to the lamellar membrane that surround the amphid/phasmid cilia, or the secreted extracellular material lining these channels. Which substrate(s) the β1,3-N-acetylglucosaminyltransferase, BGNT–1.1 (B3GNT1), glycosylates, and how this influences sensory neuron/glial cell development and function, remains to be determined in a future, detailed study of the gene.

## Conclusion

In conclusion, we demonstrated the utility and efficiency of using deep-sequenced multimutant strains in combination with SKAT to rapidly uncover novel genes required for a biological process of interest—here, ciliated sensory neuron development and/or function. The role of BGNT–1.1 in this process, seemingly independent of dystroglycan, supports the notion that Walker-Warburg syndrome may result at least in part from ciliary dysfunction, and thus could be considered a novel ciliopathy. Our findings also underscore the importance of identifying novel dye-filling genes, some of which might be implicated in human ciliopathies. For all new putative dye-filling genes highlighted in this study, we had no prior knowledge of their importance in ciliated sensory neuron function, and may not have (easily) uncovered them using alternative methods. Our approach therefore reduces the hurdle of traditional forward genetic methods, namely identifying the causative allele, and improves upon reverse genetics by allowing high gene/mutation coverage in a relatively small number of strains. Lastly, we propose that our approach is applicable not only for *C. elegans*, but any organism with a small genome that can be quickly sequenced and where numerous mutant strains can be isolated andphenotyped with relative ease, including *Drosophila* and *Arabidopsis*.

## Materials and Methods

### Strains and Maintenance

Worms were cultured on Nematode Growth Medium (NGM) seeded with Escherichia coli (0P50) at 20°C as described previously [42]. The following strains were obtained from the Caenorhabditis Genetics Center (University of Minnesota, Minneapolis, MN): N2 Bristol, CB4856, CH1869, CH1878 and PR813. VC2010, the wild-type reference strain used during the dye-filling screen, was derived from N2 [43]. The Million Mutation Project strains were isolated and their genomes’ sequenced by Thompson *et al.* [1]. The 480 Million Mutation Project strains used in this study are listed in **S1 Table**.

### Preparation of Transgenic Lines

For native rescue of VC20628 *bgnt–1.1(gk361915)*, 25 ng/μl of fosmid WRM065bB05 containing *bgnt–1.1* was injected into *bgnt–1.1* mutants along with 80 ng/μl of pRF4 *rol–6(su1006dm)* as a co-injection marker. *bgnt–1.1(gk361915);* Ex[CHE-2::GFP; pRF4] was created by crossing *bgnt–1.1(gk361915)* with wild-type worms expressing Ex[CHE–2::GFP; pRF4]. The translational Psrh–220::IFT-20::tdTomato fusion was generated as described in [11], except that tdTomato was used in place of GFP. 1 μl of the PCR product was microinjected into germline of gravid worms along with a co-injection markers (pRF4 *rol–6(su1006dm)*, final concentration of 100 ng/μl). Stable lines expressing this extrachromosomal array were crossed into DM13283 *dpy–5(e907);* sIs12964[Pgrd–15::GFP; pCeh361] to create the strain MX1924 *dpy-5(e907);* Ex[Psrh–220::IFT–20::tdTomato; pRF4]; sIs12964[Pgrd–15::GFP; pCeh361]. *bgnt–1.1(gk361915)* was also introduced to this line *via* genetic crossing to create MX2236 *bgnt*-1. *1(gk361915); dpy-5(e907);* Ex[Psrh-220::IFT-20::tdTomato; pRF4]; sIs12964[Pgrd-15::GFP; pCeh361]. The BGNT-1.1::GFP recombineered fosmid construct (Construct # 6821068113870966 H08) was obtained from the TransgeneOme (https://transgeneome.mpi-cbg.de/transgeneomics/index.html). To generate a strain expressing this construct, 25 ng/μl of the BGNT-1.1::GFP recombineered fosmid was injected into N2 worms along with 4 ng/μl of Posm-5::XBX–1::tdTomato as a cilia-marker, and 80 ng/μl of pRF4 *rol–6(su1006dm)* as a co-injection marker.

### CRISPR-Cas9 Knockout Mutant Strain Generation

VC3671 *(gk3637)*, VC3674 *(gk3674)*, and VC3675 *(gk3639*) for *bgnt–1.1/F01D4.9* were generated using the CRISPR-Cas9 system as described by [32] in an N2 VC2010 background [1]. The 20 bp guide sequence for *bgnt–1.1* was designed to include a 3’GG motif, as guides with GG at the 3’ end are purported to give higher integration efficiency [44]. 500 bp homology arms (ordered as gBlocks from IDT) were designed to flank exons 1 and 2 of *bgnt–1.1.* The homology arms were inserted into a Pmyo-2::GFP-neoR-loxP disruption/deletion vector (provided by the Calarco Lab) using Gibson Assembly. The guide sequence and homology arms sequences are available in **S7 Table** and **S8 Tables**, respectively.

Of the three null mutations, *gk3637* was generated using purified Cas9 protein, while *gk3674* and *gk3639* were generated using a plasmid-encoded version of the protein. The Cas9 protein was prepared according to the procedure described in [45] Paix et al. (2015). The protein injection mix was assembled as described in [45] and used tracrRNA and *bgnt–1.1* crRNA ordered from IDT.

Putative integrants were validated by generating PCR amplicons spanning the junction between genomic DNA and the inserted cassette. Primer F01D4.9–1–L was used in conjunction with primer pMyo–2–SEC to validate the region just upstream of the putative insertion. This generated a 1075 bp product that covers genomic DNA as well as a region within the insertion. Primer F01D4.9–1–R was used in conjunction with primer NeoR-SEC to validate the region just downstream of the putative insertion. This generated a 1632 bp product that covers genomic DNA as well as a region within the insertion. Sanger sequencing of the PCR amplicons was conducted by the Nucleic Acid Protein Service Unit (NAPS, UBC).

### Identification, Mapping and Cloning of BGNT-1.1

To rough-map the dye-filling defects of Million Mutation Project strains to an arm of a chromosome we used the high-throughput SNP mapping approach created by Davis *et al.*. The following SNPs used by Davis *et al.* [46] were omitted from our analysis because the whole genome sequence data from Thompson *et al.* [1] could not safely deduce that the SNPs from parental strain subjected to mutagenesis, VC2010 (from which the Million Mutation Project strains were generated), matched those of Bristol N2 but not Hawaiin CB4856 (mapping strain): W03D8, F58D5, T01D1, Y6D1A, Y38E10A, T12B5, R10D12, F11A1, and T24C2.

### Dye-Filling Procedures

Dye-filling assays were performed using the fluorescent dye DiI (Molecular Probes; DiIC18 Vybrant DiI cell-labelling solution, diluted 1:1000 with M9 buffer). Mixed stage *C. elegans* cultures were stained for 30 minutes, and Dil uptake into the amphid and phasmid neurons was visualised using either a Zeiss fluorescent dissection scope (dye-filling screen) or spinning disc confocal microscope (WaveFX spinning disc confocal system from Quorum Technologies) using a 25X oil (N.A 0.8) objective and Hammamatsu 9100 EMCCD camera. Volocity software (PerkinElmer) was used for acquisition. The completely dye-filling defective (*dyf* mutant strain PR813 *osm-5(p813)* was used as a positive control for the dye-filling phenotype.

For the dye-filling screen, two plates of mixed-stage *C. elegans* were dye-filled for each Million Mutation Project strain, and defects were quantified by counting the number of worms exhibiting amphid and/or phasmid dye-filling defects. A worm was classified to have a dye-filling defect if: *i*) no fluorescence was observed, *ii*) fluorescence was observed to be greatly reduced (minimum of an estimated 3x fluorescence reduction compared to wild-type staining from the experiment at the same magnification and laser intensity) and/or *iii*) fluorescence staining pattern was abrogated (e.g., accumulations of fluorescence at tips of dendrites with little to no staining in cell bodies). Fifteen worms were scored from each plate. If the dye-filling of a Million Mutation Project strain appeared qualitatively dimmer than wild-type worms across both plates or if ≥ 25% of the population exhibited a dye-filling defect the assay was repeated for that strain. A Fisher’s exact test followed by p-value adjustment using false discovery rate of 5% (Benjamini-Hochberg procedure) was used to if they exhibited a significant dye-fill defect compared to wild-type (N2). This was done separately for both amphids and phasmids.

### Imaging Sensory Neurons and Cilia

For visualisation of fluorescent-tagged proteins, worms were immobilised in 1μl of 25mM levamisole and 1μl of 0.1μm diameter polystyrene microspheres (Polysciences 00876–15, 2.5% w/v suspension) on 10% agarose pads and visualised under a spinning disc confocal microscope (WaveFX spinning disc confocal system from Quorum Technologies) using a 100X oil (N.A 1.4) objective and Hammamatsu 9100 EMCCD camera. Volocity 6.3 was used to deconvolve images as well as measure ADL cilia length and distal tip of ADL cilia to distal end of amphid socket cell length. The researcher was blind while performing the quantisation of ADL cilia/dendrite phenotypes.

### Immunofluorescence

Worms were permeabilised, fixed and stained according to standard methods [47]. To mark the *cis*-Golgi, two anti-SQL–1 antibodies, one directed against the N terminus of SQL–1 and one affinity purified antibody against the C terminus of SQL-1, were used. These antibodies have been characterised previously [33]. Both were visualised with secondary goat-anti rabbit Alexa 594 (Molecular Probes, Eugene, OR; 1:800). Localisation of BGNT–1::GFP and SQL–1 was imaged using a SpinD1454 Roper/Nikon spinning disk microscope with a 100x objective.

### SKAT Analysis

We performed SKAT using the SKAT package (version 1.0.9) [2] in R (version 3.2.4). No covariates were used. Given that the MMP library was created *via* random mutagenesis of the same isogenic parental strain [1] we did not have to control for population stratification. We chose to perform SKAT using a linear regression framework to take full advantage of the proportion data we had collected, as the logistic regression framework for SKAT only allows for a dichotomous response variable. To apply a linear regression framework to our proportion data we added a small constant to all the data points for the response variable, and then log transformed them. We used probability plots to choose the best constant (S2 & S3 Fig), and thus used a constant of 0.005 for the amphid phenotype data, and a constant of 0.05 for the phasmid phenotype data.

Custom, biologically relevant weights were assigned to the variants. Nonsense, splicing mutations and frameshift causing deletions were assigned a weight of 1, in-frame deletions were assigned a weight of 0.75, and missense mutations were assigned a weight of 0.25. Gene-based tests for all genes with a minor allele count > 6 were performed. A false discovery rate (Benjamini-Hochberg procedure) of 5% was used to determine genes which were significantly associated with the phenotype. Make, Bash, Perl and R scripts used to perform the analysis can be found at:https://github.com/ttimbers/Million-Mutation-Project-dye-filling-SKAT.git

### Power Analysis

To estimate power and recommend a minimum sample size for future experiments we performed a bootstrap power analysis using the amphid dataset. To do this, we randomly sampled (resampling = FALSE) N strains from the dataset we collected, and performed the SKAT analysis presented in this paper. We did this 100 times for N = 50, 100, 200, 300 and 400. We then estimated power as the proportion of times we observed a gene to be significantly associated with the phenotype. This was also done for two, three, four and five genes. The code used to perform this analysis can also be found in the Github repository for this study: https://github.com/ttimbers/Million-Mutation-Project-dye-filling-SKAT.git

### Phylogenetic Analysis

Protein sequences (obtained from: http://www.cazy.org/) were aligned using MUSCLE 3.7 [48]. The phylogenetic tree was built using PhyML 3.0 aLRT [49] and viewed using FigTreeersion 1.3.1 (http://tree.bio.ed.ac.uk/software/figtree/).

## Acknowledgements

T.A.T. acknowledges a Banting Postdoctoral Fellowship from the Canadian Institutes of Health Research (CIHR). M.R.L. acknowledges funding from CIHR (MOP82870) and a senior scholar award from Michael Smith Foundation for Health Research (MSFHR). The work in the D.G.M. laboratory was supported by CIHR. D.G.M. is a Senior Fellow of the Canadian Institute For Advanced Research. S.J.G. acknowledges funding from a Natural Sciences and Engineering Research Council (NSERC) Undergraduate Student Research Award (USRA).

## Supplementary Information Captions

**S1 Fig.**
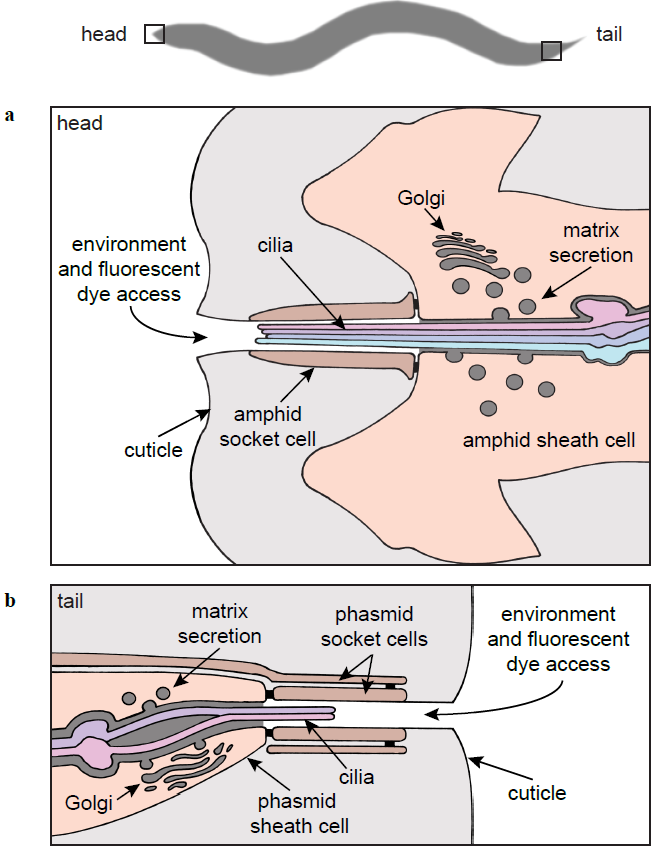
**Schematic of a longitudinal section through the wild-type amphid and phasmid sensillum.** (A) The socket and sheath cells form the amphid channel through which the cilia are exposed to the environment. The lipophilic dye, DiI, is also presumed to access the cilia *via* this channel. Cuticle, continuous with the external surface of the worm, lines the socket channel. Matrix is secreted by the Golgi apparatus of the sheath cell, and fills the space that surrounds the cilia. (B) The phasmid sensillum is organised in the same manner as described for the amphid sensillum.

**S2 Fig.**
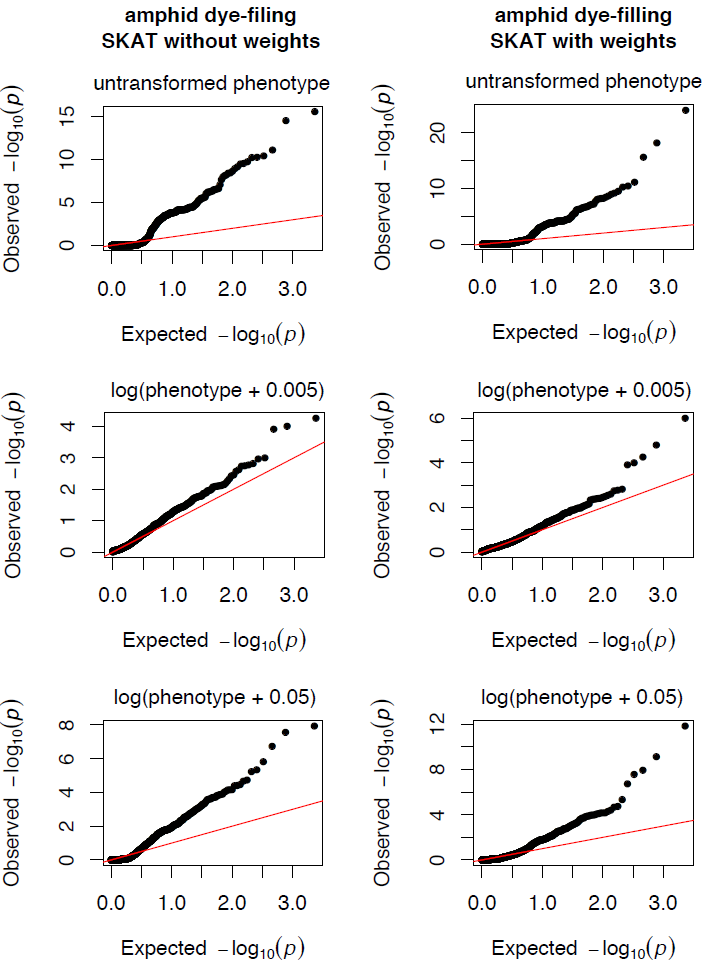
**QQ-plots of the distribution of p-values calculated with SKAT (with and without weights) using the amphid dye-filing dataset.** From these plots we chose to log transform the amphid phenotype data (proportion of dye-filling defects) and to add a constant of 0.005 to all values prior to log transformation.

**S3 Fig.**
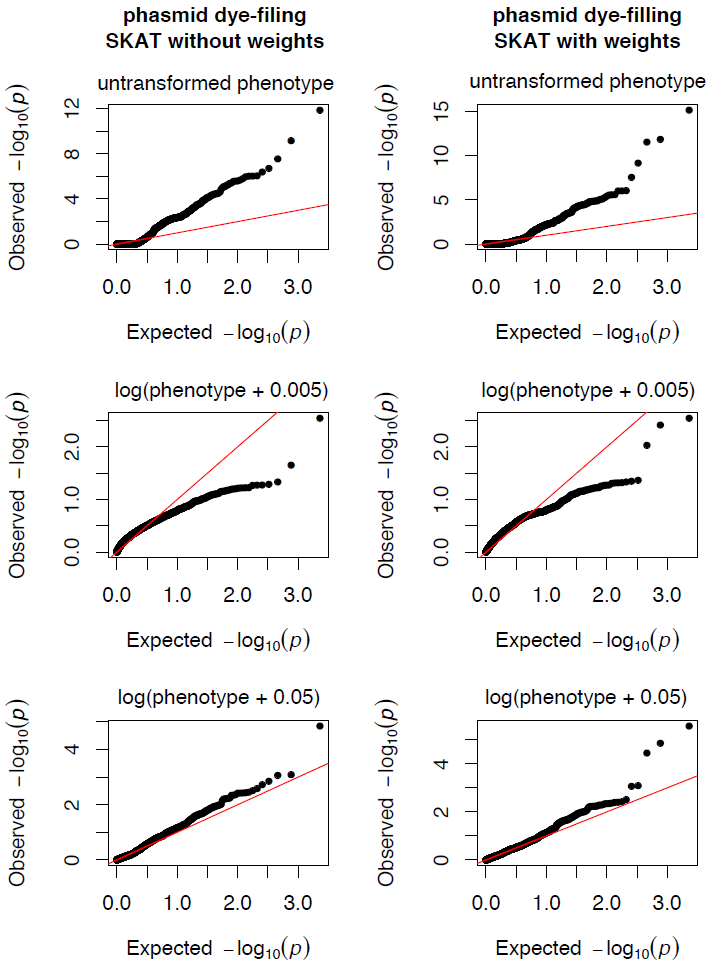
**QQ-plots of the distribution of p-values calculated with SKAT (with and without weights) using the phasmid dye-filing dataset.** From these plots we chose to log transform the phasmid phenotype data (proportion of dye-filling defects) and to add a constant of 0.05 to all values prior to log transformation.

**S4 Fig.**
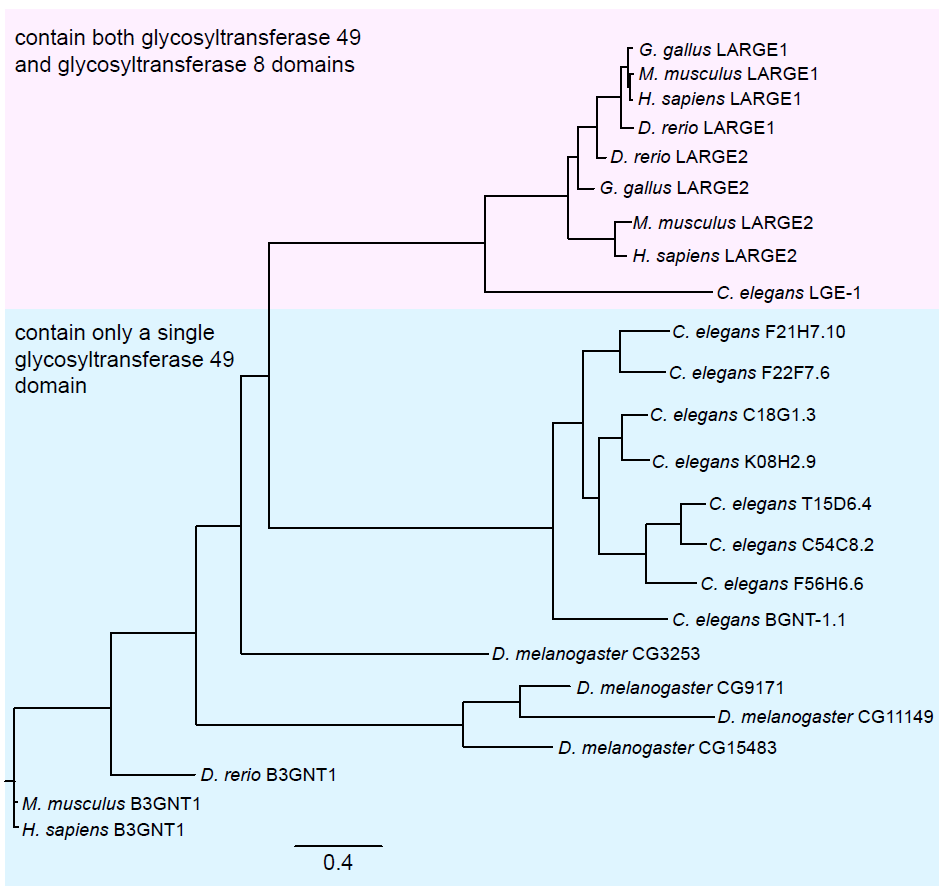
**Phylogenetic tree of glycosyltransferase 49 domain containing proteins.** The glycosyltransferase 49 domain containing proteins appear to have radiated in *C. elegans.* BGNT–1.1 is the most basal member of this family, and most closely related to vertebrate B3GNT1/ B4GAT1. Protein sequences (obtained from: (http://www.cazy.org/) were aligned using MUSCLE 3.7 (Edgar, 2004), and the phylogenetic tree was built using PhyML 3.0 aLRT (Guindon et al., 2010) and viewed using FigTree version 1.3.1 http://tree.bio.ed.ac.uk/software/figtree/).

**S5 Fig.**
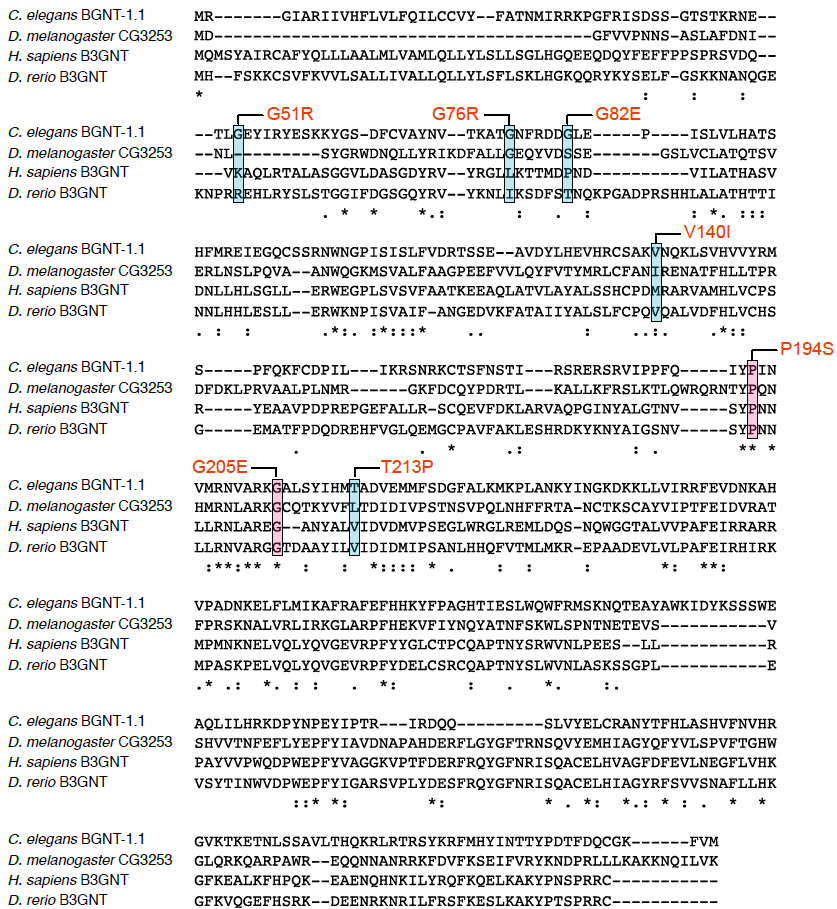
**Multiple sequence alignment of *C. elegans* BGNT–1.1 with homologous glycosyltransferase 49 domain containing proteins.** In VC20615 the *bgnt–1(gk355974)* mutation results in P194S at a conserved P (highlighted in pink). In VC20628 *the bgnt–1(gk361915)* mutation results in G205E at a conserved E (highlighted in blue). Alignment was performed by L-INS-i MAFFT using BLOSUM62 as a scoring matrix and a gap onset penalty of 1.53.

**S6 Fig.**
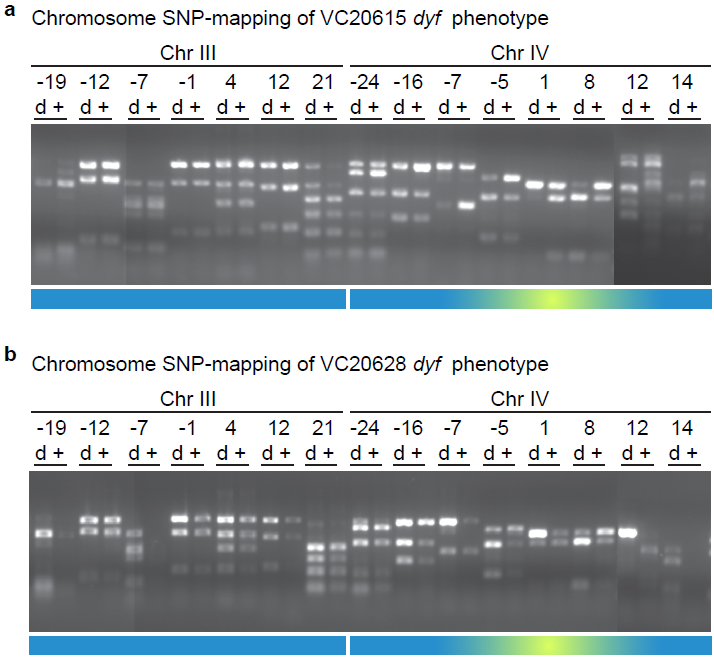
**VC20615 and VC20628 dye-fill defects map to the *bgnt-1* locus.** (A) Chromosome SNP mapping of VC20615's dyf phenoype. Each pair of lanes shows results from the SNP at the indicated genetic map position, using DNA template from either worms exhibiting dyf (d) or wild-type (+) dye-filling. Linkage is visible as an increase in the proportion of Bristol N2 DNA in dyf lanes compared to the wild-type lanes, and is visible on ChrIV from −5 to 8. (B) chromosome mapping of VC20628’s dyf phenoype. Similar to VC20615, linkage is visible as an increase in the proportion of Bristol N2 DNA in dyf lanes compared to the wild-type lanes, and is visible on ChrIV from ȡ5 to 8. All PCR samples from SNP-mapping of each strain were run on the same large gel from which multiple images were captured to visualize all samples.

**S7 Fig.**
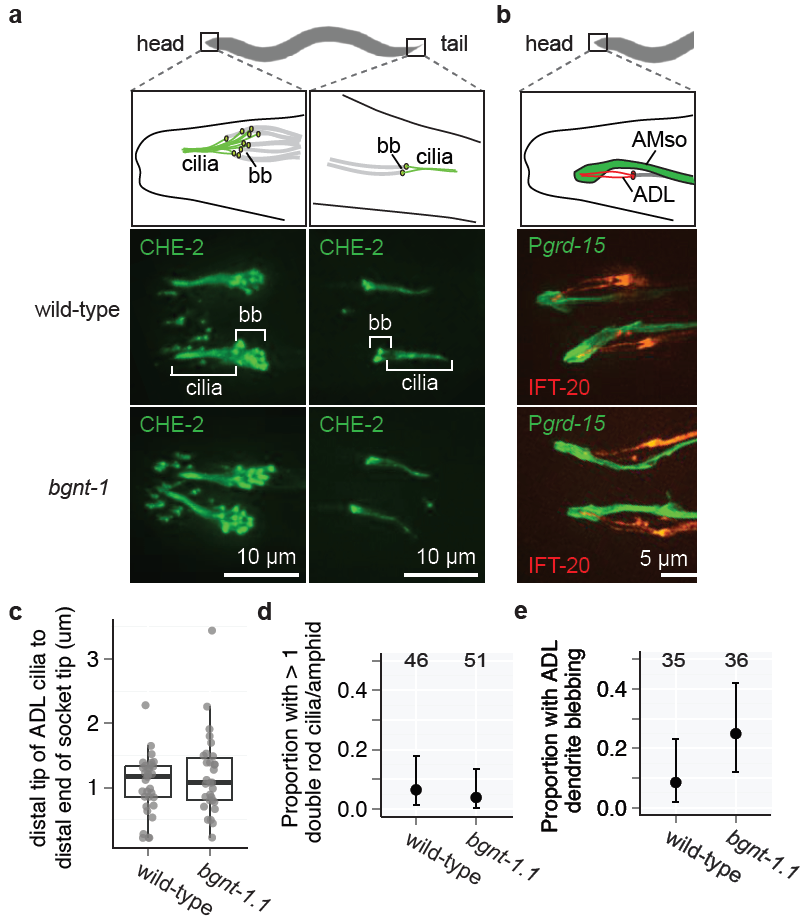
**Ciliary and socket cell structures in the *bgnt-1* mutant are present and appear superficially wild-type**. (A) Both amphid and phasmid cilia appear superficially wild-type. GFP-tagged CHE–2 (mammalian IFT80 orthologue) is used as a pan-cilia marker which localises to the basal bodies (bb) and axonemes. (B) ADL cilia correctly enter the amphid socket (AMso) cells in *bgnt–1.1* mutants. ADL cilia are labelled with Psrh–220::IFT–20::tdTomato. IFT–20 (IFT20) localises to cilia basal bodies (bb) and axonemes. The *srh–220* promoter drives expression primarily in ADL neurons. The amphid socket (AMso) cells are labelled with cytoplasmic GFP driven by an amphid socket specific promoter, *grd-15* (Hunt-Newbury et al. 2007). (C) ADL cilia penetrate the sockets cells to an equivalent depth in wild-type and *bgnt–1.1* mutants (p > 0.05, Kruskal-Wallis test). (D) ADL cilia/dendrites exhibit no guidance defects in *bgnt–1.1* mutants as assessed by the number of double-rod cilia observed in each amphid when ADL was driven by the primarily ADL specific *srh-220* promoter (p > 0.05, Fisher's exact test). Error bars represent 95% confidence intervals (Pearson Clopper method). (E) *bgnt–1.1* mutants do not exhibit a significant increase in the proportion of ADL neurons with dendritic blebbing (p > 0.05, Fisher’s exact test). Error bars represent 95% confidence intervals (Pearson Clopper method).

**S8 Fig.**
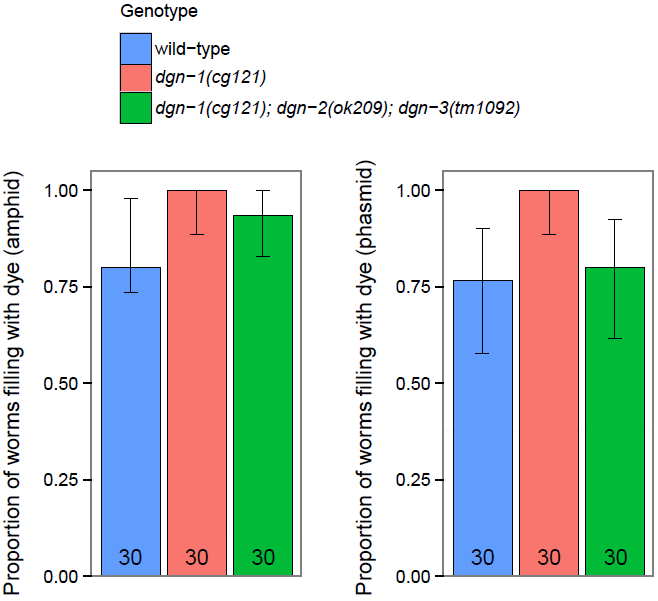
***C. elegans* Dystroglycan homologues, *dgn-1, dgn-2* and *dgn-3*, are not required for dye-filling of amphid or phasmid ciliated sensory neurons**. *dgn* mutants did not exhibit significant dye-filling defects when compared to wild-type (p > 0.05, Fisher's exact test followed by a Bonferroni correction) Error bars represent 95% confidence intervals (Clopper Pearson method).

**S9 Fig.**
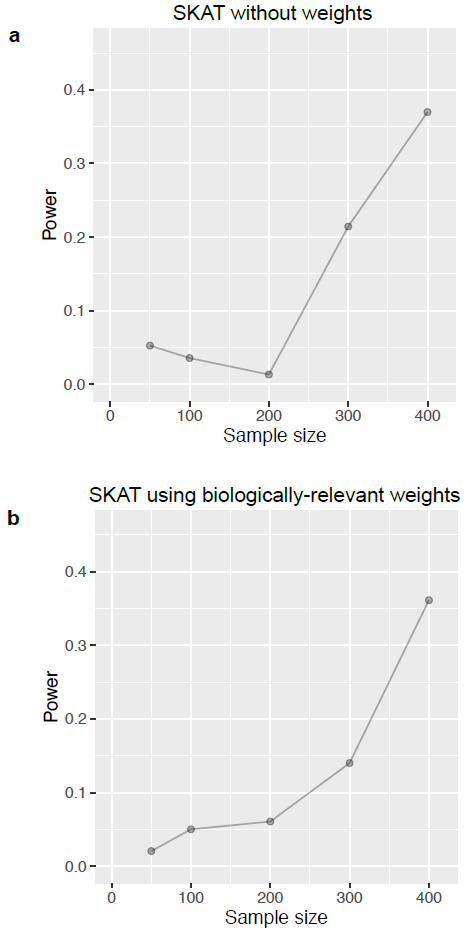
**Power analysis of SKAT analysis of amphid dye-filling data**. Raw amphid dye-filling phenotype and genotype data was randomly sub-sampled (without replacement) and analysis was performed via SKAT with (B), and without (A), biologically relevant weights. This was done 100 times for each sample size (50, 100, 200, 300, 400). For each sample size, power was calculated as the proportion of times the analysis found a gene to be significantly associated with the phenotype.

**S10 Fig.**
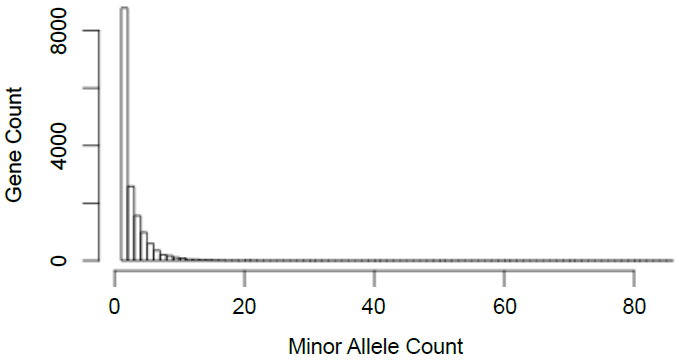
**Distribution of variant counts for mutated genes from the 480 strains screened in this study from the Million Mutation project**. The majority of genes have only a single mutation in the 480 strains screened.

**S1 Table.MMP strains phenotyped for dye-filling defects and their respective phenotypes**.

**S2 Table.Strains which exhibit a dye-fill defect and the known *dyf* genes which are mutated in these strains**

**S3 Table.SKAT analysis of amphid dye-filling phenotype using biologically relevant weights.**

**S4 Table.SKAT analysis of phasmid dye-filling phenotype using biologically relevant weights.**

**S5 Table.SKAT analysis of amphid dye-filling phenotype where all variants were weighted equally.**

**S6 Table.SKAT analysis of phasmid dye-filling phenotype where all variants were weighted equally**.

